# A novel glideosome-associated protein S14 coordinates sporozoite gliding motility and infectivity in mosquito and mammalian hosts

**DOI:** 10.1101/2023.08.30.555469

**Authors:** Ankit Ghosh, Aastha Varshney, Sunil Kumar Narwal, Nirdosh, Sachin Gaurav, Roshni Gupta, Shakil Ahmed, Satish Mishra

## Abstract

*Plasmodium* sporozoites are the infective forms of the malaria parasite in the vertebrate host. Gliding motility allows sporozoites to migrate and invade the salivary gland and hepatocytes. Invasion is powered by an actin-myosin motor complex linked to glideosome. However, the gliding complex and the role of several glideosome-associated proteins (GAPs) are poorly understood. In silico analysis of a novel protein, S14, which is uniquely upregulated in salivary gland sporozoites, suggested its association with glideosome-associated proteins. We confirmed *S14* expression in sporozoites using real-time PCR. Further, the *S14* gene was endogenously tagged with 3XHA-mCherry to study expression and localization. We found its expression and localization on the inner membrane of sporozoites. By targeted gene deletion, we demonstrate that S14 is essential for sporozoite gliding motility, salivary gland, and hepatocyte invasion. The gliding and invasion-deficient *S14* KO sporozoites showed normal expression and organization of IMC and surface proteins. Using in silico and the yeast two-hybrid system, we showed the interaction of S14 with the glideosome-associated proteins GAP45 and MTIP. Together, our data show that S14 is a glideosome-associated protein and plays an essential role in sporozoite gliding motility, which is critical for the invasion of the salivary gland, hepatocyte, and malaria transmission.

## Introduction

The *Plasmodium* life cycle alternates between a mosquito and a mammalian host, which involves invasive and replicative stages. When a mosquito probes for the blood meal in an infected human, it ingests the gametocytes, which develop further into gametes that fuse to form the zygote. The zygote transforms into ookinetes, producing hundreds to thousands of oocysts in the mosquito midgut. Following oocyst rupture, sporozoites are released in the hemolymph and further invade the salivary gland (Klug & Frischknecht, 2017; Douglas *et al*, 2015). Sporozoites transform into liver stages after transmission to the human host, forming thousands of merozoites that initiate the erythrocytic cycle (Ripp *et al*, 2021; Prudêncio *et al*, 2006). Sporozoites are highly motile cells and rely on gliding motility to travel through different host species. The gliding motility is powered by the inner membrane complex (IMC) anchored actomyosin machinery termed the glideosome (Baum *et al*, 2008; Kono *et al*, 2013). The motor is located in the IMC, separating the flattened alveolar sacs and parasite plasma membrane (PPM) (Gould *et al*, 2011). The gliding-associated proteins (GAPs) tether MyoA to the IMC and hold the PPM and the IMC together (Boucher & Bosch, 2015). These transmembrane proteins connect the submembrane motor to the extracellular environment (King, 1988)

*Plasmodium* sporozoites can invade target cells in the mosquito and the mammalian hosts and many proteins that have been implicated during this process are expressed specifically in sporozoites. Thrombospondin-related anonymous protein (TRAP) was found to be the first protein involved in sporozoite gliding motility and host cell invasion. TRAP mutant sporozoites failed to invade the salivary gland, and hemolymph sporozoites were nonmotile (Sultan *et al*, 1997). TRAP cytoplasmic tails incorporate a C-terminal tryptophan residue that is crucial for interaction with aldolase that connects with an actin-myosin-based motor (Heintzelman, 2015; Buscaglia *et al*, 2003). S6 is also a TRAP family adhesion, and by disruption, its role has been implicated in parasite adhesion and gliding motility (Steinbuechel & Matuschewski, 2009). The other proteins involved in parasite motility and host cell invasion include MAEBL (Kariu *et al*, 2002; Saenz *et al*, 2008), TLP (Heiss *et al*, 2008), the rhoptry-resident proteins TRSP (Labaied *et al*, 2007), RON4 (Giovannini *et al*, 2011), GEST (Talman *et al*, 2011), TREP/UOS3 (Mikolajczak *et al*, 2008; Combe *et al*, 2009), the GPI-anchored circumsporozoite protein (CSP) of the sporozoite and the small solute transporter PAT (Kehrer *et al*, 2016) and claudin-like apicomplexan microneme protein (CLAMP) (Loubens *et al*, 2023). The individual functions of these proteins are known. However, how they interact with each other to coordinate gliding motility and invasion is poorly understood.

These proteins are specific to sporozoites; however, several overlapping proteins are known that function in both merozoites and sporozoites. Proteins such as GAP-40, −45, and −50 together with myosin A tail domain interacting protein (MTIP) cluster MyoA with IMC (Poulin *et al*, 2013; Daher & Soldati-Favre, 2009). In *P. falciparum*, the peripheral protein GAP45 is myristoylated and palmitoylated, possibly required for membrane targeting (Rees-Channer *et al*, 2006). GAP45 in *Toxoplasma gondii* is involved in the recruitment of the motor complex (Frénal *et al*, 2010). *Pf*GAP50 is a transmembrane protein that anchors the invasion machinery in the inner membrane complex (Baum *et al*, 2006; Yeoman *et al*, 2011). In *T. gondii*, GAP45 and GAP50 form a complex with MyoA and its light chain, MLC1.

While the glideosome and many surface proteins have been studied and found essential for gliding motility and invasion, the role of gliding-associated proteins is unexplored. To identify a novel GAP, we performed bioinformatic studies on S14, whose transcripts were highly upregulated in sporozoites in a suppressive subtraction hybridization study (Kaiser *et al*, 2004). Similar to GAP45 and −50, we found that S14, while not containing transmembrane domains or signal peptides, is predicted to be secreted via the nonclassical pathway. It also contains a predicted palmitoylation signal, possibly indicating its membrane targeting (Table S1). In this study, we investigated the role of S14 in the *P. berghei* life cycle. We demonstrate that S14 is an IMC protein interacting with the glideosome-associated proteins MTIP and GAP45. We disrupted the gene and found that S14 is essential for sporozoite gliding motility, host cell invasion, and malaria transmission.

## Results

### In silico studies show that S14 is secreted via a nonclassical pathway

A study comparing transcriptome differences between sporozoites and merozoites using suppressive subtraction hybridization found several genes highly upregulated in sporozoites and named them ‘S’ genes (Kaiser *et al*, 2004). We narrowed it down to a candidate named S14, which lacked signal peptide and transmembrane domains. In silico studies revealed that *Pb*S14 is a protein conserved in all *Plasmodium* species and has no similarity in other organisms (Figure S1). The identity matrix showed that the *Pb*S14 protein is highly conserved among all Plasmodia (Table S2). To understand its possible function, we analyzed several sporozoite-specific proteins. We found that gliding-associated proteins GAP45 and GAP50 show similar properties and contain a palmitoylation signal, secreted via the nonclassical pathway and localized to the inner membrane complex (Rees-Channer *et al*, 2006). Next, we performed bioinformatic analysis to check whether S14 is targeted to the inner membrane by a nonclassical secretion pathway. We analyzed the presence of transmembrane domains, prediction of palmitoylation signals, and interactions as previously described (Boucher and Bosch, 2015). We found that S14, while not containing transmembrane domains or signal peptides, is predicted to be secreted via the nonclassical pathway. It also contains a predicted palmitoylation signal, possibly indicating its membrane targeting (Table S1). The nonclassical secretion pathway, which depends on palmitoylation and myristoylation, has been identified in most eukaryotes (Rabouille et al. 2012), including *P. falciparum* (Moskes et al. 2004).

### S14 is expressed and localized on the membrane of sporozoites

We started our study by validating the transcripts of S14 in different stages of the parasites using quantitative real-time PCR. We found that S14 was predominantly expressed in midgut and salivary gland sporozoites (Figure 1A). To further investigate whether S14 transcripts are translated, we endogenously tagged the *S14* gene with 3XHA-mCherry using double crossover homologous recombination (Figure S2A). The correct site-specific integration was confirmed by diagnostic PCR (Figure S2B). We initiated the mosquito cycle and found that the C-terminal tag did not affect parasite development throughout the life cycle stages (Figure S2C and S2D). We monitored live mCherry expression throughout the parasite life cycle stages. S14-3XHA-mCherry expression was observed in sporozoites but not in the blood and liver stages (Figure 1B and C). Analysis of the mCherry pattern on sporozoites revealed localization on the membrane of the sporozoites (Figure 1D). Furthermore, IFA with anti-CSP and anti-mCherry antibodies confirmed S14 localization on the membrane (Figure 1E). Expression of the S14-3XHA-mCherry fusion protein (∼66.5 kDa) was also confirmed by Western blotting using an anti-mCherry antibody. The appearance of an extra band with higher molecular weight in immunoblotting was possibly due to the palmitoylation of S14 (Figure 1F). Similar electromobility shifts of GAP45 due to the palmitoylation have been reported (Frénal *et al*, 2010). These results indicate that S14 is a sporozoite-specific membrane protein.

**Figure 1.**
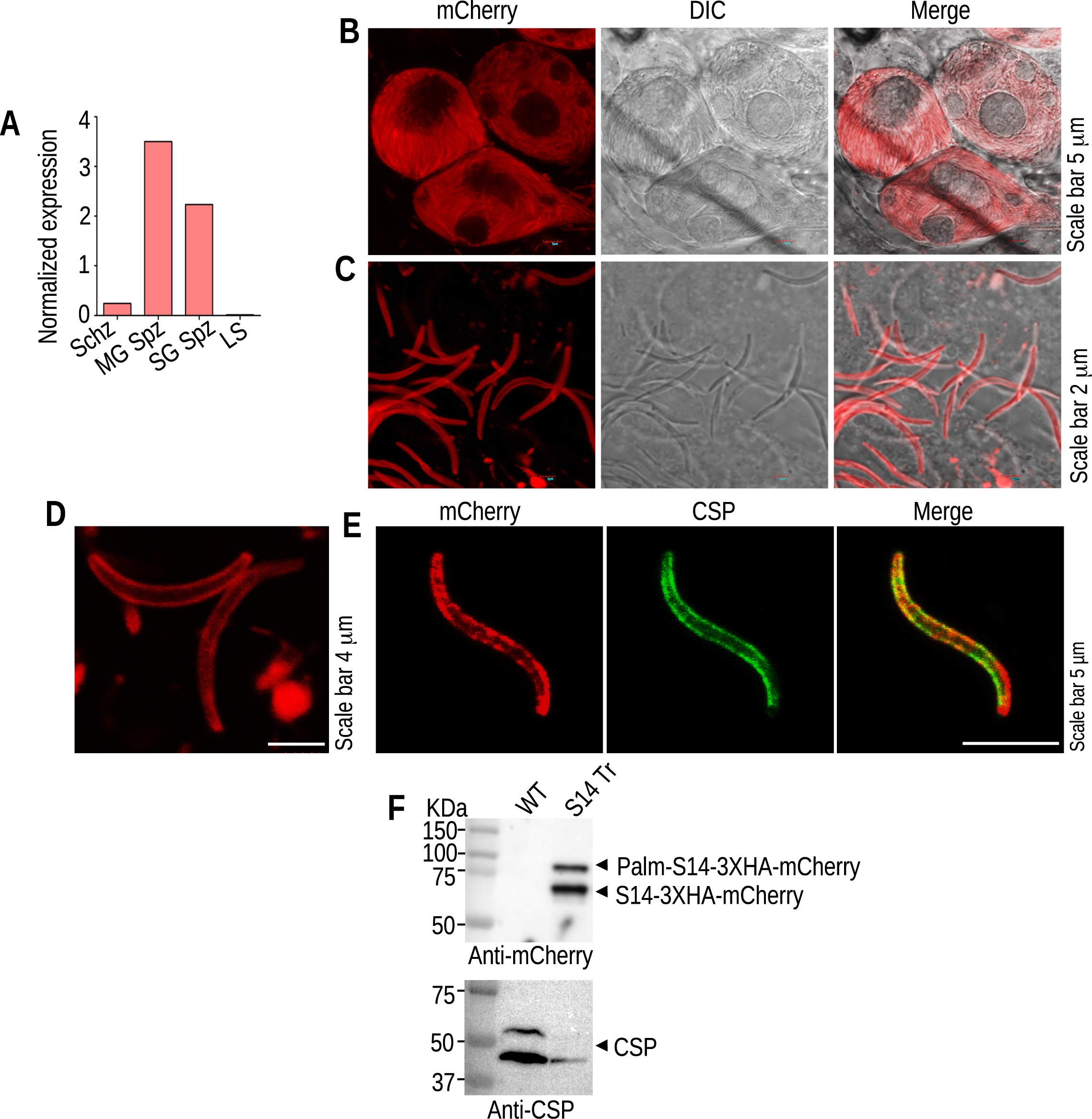
*S14* expression and localization. **(A)** The gene expression of *S14* was analyzed using quantitative real-time PCR, which revealed the highest expression in sporozoites. The expression of *S14* was normalized to the *PbHsp70* transcript. BS; blood stages, Schz; schizonts, MG Spz; midgut sporozoites, SG Spz; salivary gland sporozoites, LS; liver stages. **(B)** Live microscopy image of midgut oocysts expressing the mCherry reporter. **(C)** mCherry-expressing salivary gland sporozoites. **(D)** Salivary gland sporozoites expressing S14-3XHA-mCherry on the membrane. **(E)** Confirmation of S14 expression on the membrane of the sporozoites by coimmunostaining with surface protein marker anti-CSP antibody. **(F)** Western blot analysis of S14-3XHA-mCherry salivary gland sporozoite lysates. The blot was first probed with an anti-mCherry antibody, then stripped and reprobed with an anti-CSP antibody.

### *S14* KO sporozoites egress from the oocyst normally while failing to invade the salivary gland

To investigate the role of S14 in the *P. berghei* life cycle, we disrupted the gene by double-crossover homologous recombination (Figure S3A). The drug-resistant parasites expressing GFP indicated successful transfection (Figure S3B). We obtained clonal lines of the KO parasites by limiting dilution of the parasites, integration, and the absence of WT locus was confirmed by diagnostic PCR (Figure S3C). Finally, two *S14* KO clonal lines were confirmed by Southern blotting, which showed the modified locus in the KO parasites (Figure S3D). The S14-complemented parasite line was also generated to check the specificity of the phenotype (Figure S3E). First, a new *S14* KO parasite with yFCU was generated, and then an S14 expression cassette consisting of the 5’UTR, ORF, and 3’UTR was amplified and transfected into *S14* KO (yFCU) parasites. After recombination, the WT locus was amplified in S14 comp parasites (Figure S3F). Next, we checked the propagation of the asexual intraerythrocytic cycle of KO parasites, which was comparable to that of WT GFP parasites (Figure S4). To analyze the phenotype of *S14* KO parasites in mosquito stages, we transmitted them to mosquitoes by allowing them to probe for a blood meal. We observed the mosquito midgut and salivary glands on day 14 and day 19 post-blood meal. We found that oocyst formation and development of sporozoites were comparable to those of WT GFP parasites (Figure 2A-D). However, no sporozoite-associated GFP signals were observed in salivary glands, and the number of salivary gland sporozoites per mosquito was severely reduced (Figure 2E and F). Genetic complementation of the KO parasites restored salivary gland sporozoite numbers to a level similar to WT GFP parasites (Figure 2F). To investigate whether the KO sporozoites failed to egress from the oocyst or could not invade the salivary gland. We counted the sporozoite numbers in the mosquito hemolymph. We found a higher accumulation of hemolymph sporozoites in KO-infected mosquitoes than in WT GFP-infected mosquitoes (Figure 2G), suggesting that mutant sporozoites failed to invade the salivary gland. These results indicate that mutant sporozoites egress from the oocysts normally but fail to invade salivary glands.

**Figure 2.**
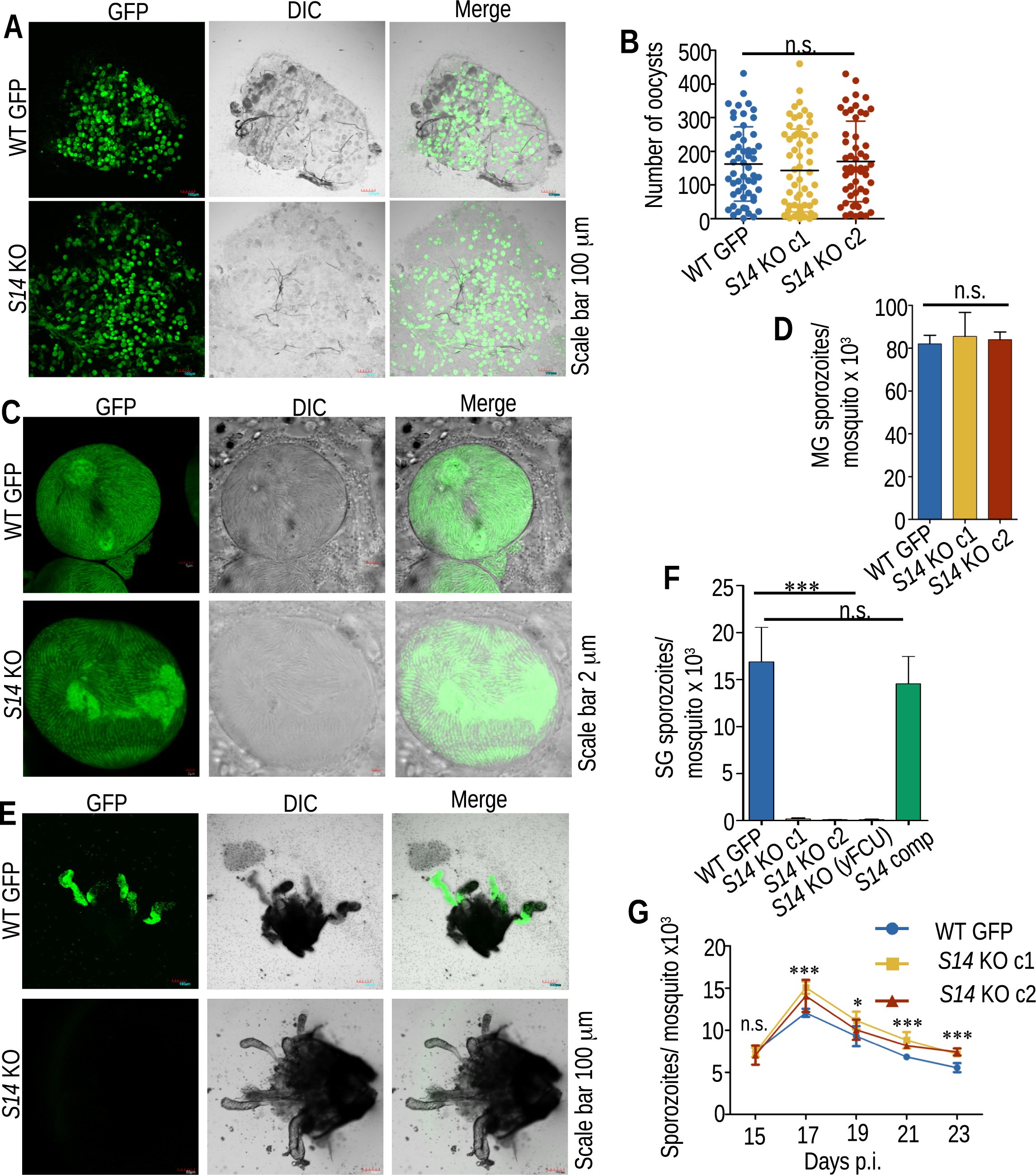
S14 is essential for salivary gland invasion by sporozoites. **(A)** Live microscopy images of the mosquito midgut showing oocysts. **(B)** The number of oocysts in WT GFP and *S14* KO parasites was not significantly different (P= 0.952, one-way ANOVA). **(C)** Live microscopy images of sporulating oocysts of WT GFP and *S14* KO parasites. **(D)** Midgut oocyst sporozoite number, no significant difference (P= 0.943, one-way ANOVA). **(E)** Dissected salivary glands showed GFP-expressing sporozoites in WT GFP, while no GFP-expressing sporozoites were observed in *S14* KO. **(F)** Salivary gland sporozoite numbers, negligible sporozoites in *S14* KO, a significant difference (***P<0.0008, one-way ANOVA). Complementation restored the sporozoites number in *S14* KO parasites at the WT GFP levels, no significant difference (P= 0.434, Student’s t test) **(G)** *S14* KO and WT GFP hemolymph sporozoites were collected on the indicated days post blood meal, and sporozoite numbers were quantified. We found a comparable number of sporozoites on day 15, with no significant difference (P=0.621). There was a higher accumulation of hemolymph sporozoites in S14 KO on days 17-23 with a significant difference from WT GFP (day 19 *P=0.0187, day 17 ***P<0.0001, days 21 and 23 ***P=0.0006, one-way ANOVA).

### S14 is essential for malaria transmission

To evaluate the ability of salivary gland sporozoites to transmit malaria, infected mosquitos were allowed to inoculate sporozoites in C57BL/6 mice by enabling them to probe for the blood meal. We found that mice inoculated with WT GFP or S14 comp sporozoites became patent on day 3, whereas *S14* KO sporozoites failed to initiate blood-stage infection (Table 1). Next, we checked whether this in vivo infectivity defect was due to negligible salivary gland sporozoite load or mutant sporozoites failed to infect mice. For this, we intravenously inoculated C57BL/6 mice with 5,000 hemolymph sporozoites and found that all the mice inoculated with KO sporozoites failed again to initiate blood-stage infection (Table 1). To determine the stage-specific defect, we performed in vitro assays. First, we infected HepG2 cells with hemolymph sporozoites and observed EEF development at 40 hpi. In three independent experiments, we found EEFs in culture inoculated with WT GFP sporozoites but not in the KO-infected cells (Figure 3A and S5A). Next, we checked the invasion capacity of *S14* KO hemolymph sporozoites using double immunostaining, which differentiates outside vs inside sporozoites (Rénia *et al*, 1988). *S14* KO sporozoites showed a complete failure to invade hepatocytes (Figure 3B and S5B), explaining the inability to establish an infection in mice. Next, we performed a development assay to check the mutant sporozoite transformation ability into a bulb-like structure. For this, we incubated the mutant sporozoites in a transformation medium for 4 h. We found that *S14* KO sporozoites retain the ability to transform into bulb-like structures (Figure 3C and D). These data demonstrate that *S14* KO sporozoites lost infectivity to mammalian hosts due to the inability of sporozoites to invade cells.

**Figure 3.**
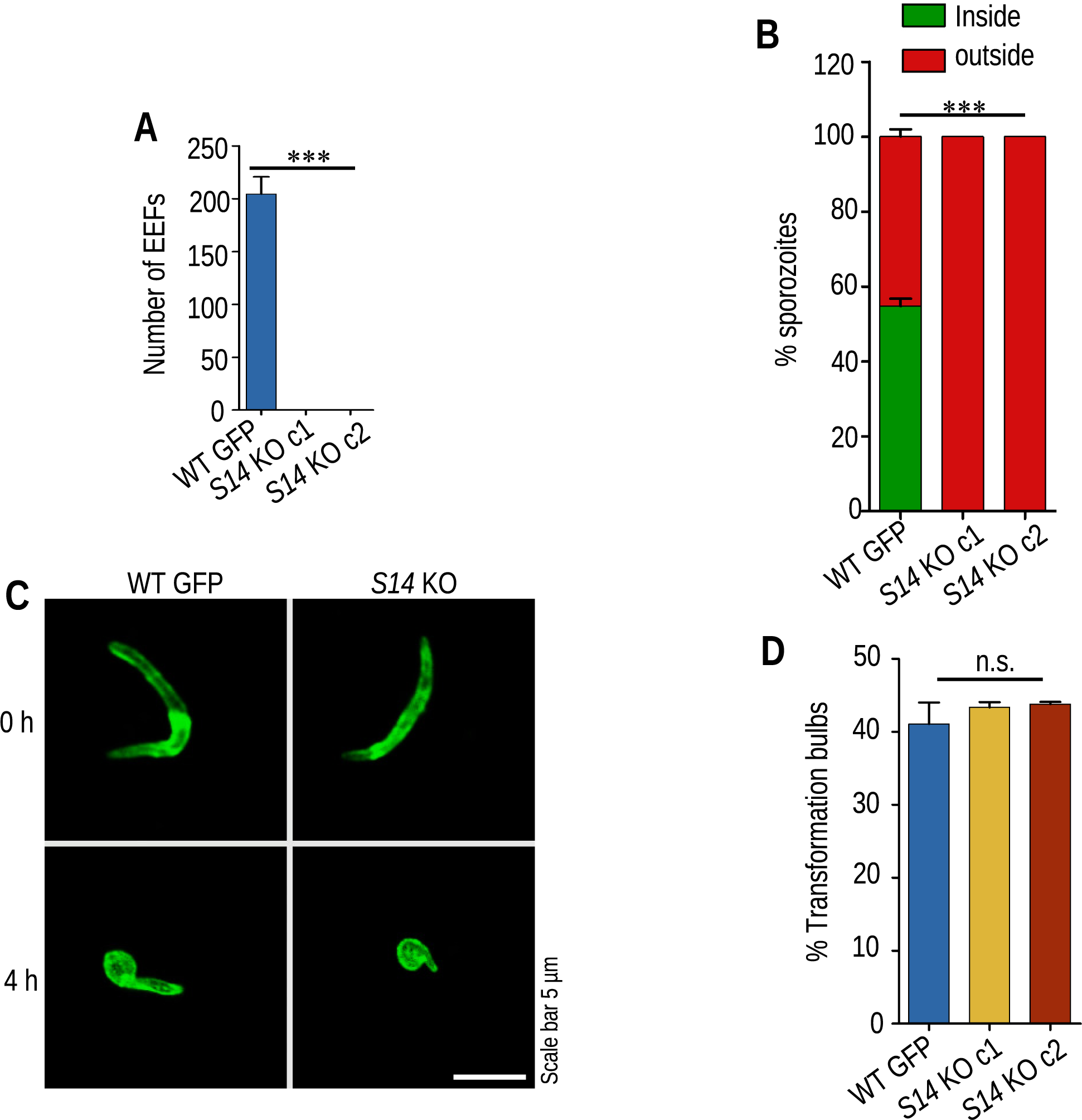
S14 is essential for hepatocyte invasion and malaria transmission. **(A)** HepG2 cells infected with hemolymph sporozoites were immunostained, and EEF numbers were quantified. No EEFs were observed in *S14* KO parasites, a significant difference (***P<0.0001, one-way ANOVA). **(B)** Quantification of sporozoites inside vs outside in invasion assay. All sporozoites were found to be outside in *S14* KO parasites, a significant difference (***P<0.0001, one-way ANOVA). **(C)** *S14* KO and WT GFP hemolymph sporozoites transformed into bulbs after incubation for 4 h in an activation medium. **(D)** Quantifying sporozoite transformation into bulbs, no significant difference (P=0.419, one-way ANOVA).

**Table 1.** Infectivity of *S14* KO sporozoites in C57BL/6 mice. C57BL/6 mice were inoculated with WT GFP, *S14* KO or *S14* comp sporozoites by mosquito bite or intravenous injection. The appearance of parasites in the blood was confirmed by making Giemsa-stained blood smear.

| Experiment | Parasites | Route | Number of sporozoites injected | Mice positive/Mice infected | Pre-patent period (days) |
| --- | --- | --- | --- | --- | --- |
| 1 | WT GFP | Mosquito bite | From 10 mosquitos | 4/4 | 3 |
|  | *S14* KO c1 | Mosquito bite | From 10 mosquitos | 0/4 | NA |
|  | *S14* KO c2 | Mosquito bite | From 10 mosquitos | 0/4 | NA |
| 2 | WT GFP | intravenous | 5,000 | 5/5 | 3 |
|  | *S14* comp | intravenous | 5,000 | 5/5 | 3 |
|  | *S14* KO c1 | intravenous | 5,000 | 0/5 | NA |
|  | *S14* KO c2 | intravenous | 5,000 | 0/5 | NA |
|  | *S14* KO (yFCU) | intravenous | 5,000 | 0/5 | NA |
| 3 | WT GFP | intravenous | 5,000 | 0/4 | 3 |
|  | *S14* KO c1 | intravenous | 5,000 | 0/4 | NA |
|  | *S14* KO c2 | intravenous | 5,000 | 0/4 | NA |

### S14 localizes on the inner membrane and is essential for parasite gliding motility

Invasion-deficient *S14* KO sporozoites indicate that either S14 interacts with host receptors or cannot generate power to invade cells. As S14 lacks a signal sequence and transmembrane domain, we analyzed whether S14 is present on the outer or inner membrane. We treated S14-3XHA-mCherry hemolymph sporozoites with Triton X-100 to remove the outer membrane. The Triton X-100-treated and untreated sporozoites were immunostained with anti-CSP and anti-mCherry antibodies. The CSP signal was lost in Triton X-100-treated sporozoites, whereas the S14-3XHA-mCherry signal was retained (Figure 4A). Western blotting confirmed the IFA result, which detected the mCherry signal in Triton X-100-treated sporozoites (Figure 4B). To further confirm the inner membrane localization of S14, we generated antibodies against two IMC proteins MTIP (Bergman *et al*, 2003) and GAP45 (Gaskins *et al*, 2004), and performed IFA. The MTIP and GAP45 signals were retained in Triton X-100-treated sporozoites and colocalized with the mCherry signal (Figure 4A). This result indicates that S14 is present within the inner membrane of sporozoites. *Plasmodium* parasites actively invade host cells, powered by gliding motility (Frénal *et al*, 2017). Next, we checked the gliding motility of WT GFP and *S14* KO hemolymph sporozoites. WT GFP sporozoites glided normally, whereas *S14* KO sporozoites were found to be nonmotile (Figure 4C). We counted the sporozoites associated with or without trails to quantify the percentage of gliding sporozoites. Approximately 53% of WT GFP sporozoites were associated with trails, whereas no trails associated with sporozoites were observed in *S14* KO (Figure 4D). These data demonstrate that S14 is an IMC protein that powers the sporozoite’s gliding motility.

**Figure 4.**
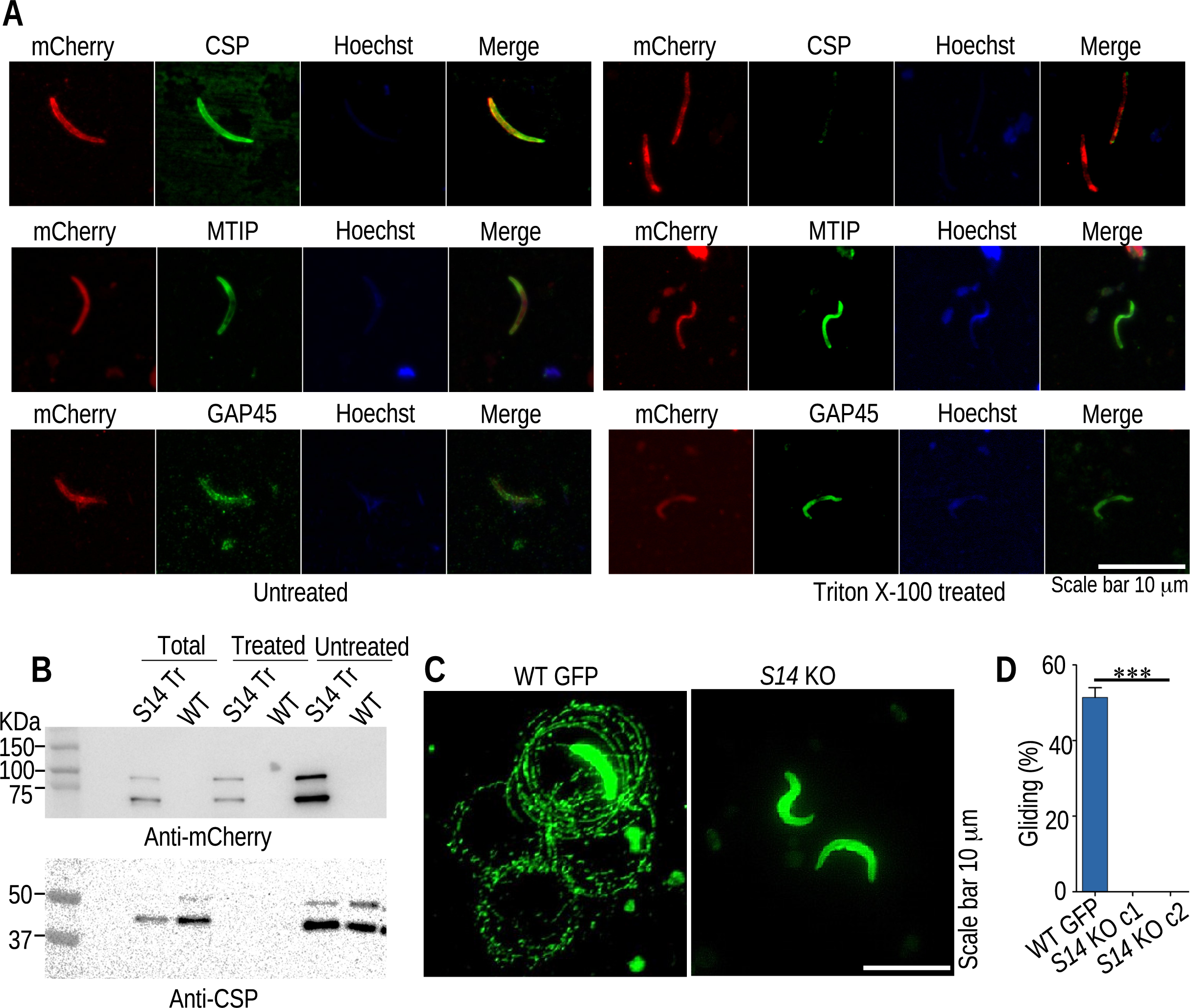
S14 is an inner membrane protein and is essential for parasite gliding motility. **(A)** Triton X-100-treated sporozoites retained mCherry staining, which colocalized with MTIP and GAP45, while CSP staining was lost, suggesting that S14 is present within the inner membrane of the sporozoites. **(B)** Triton X-100-treated and untreated sporozoites were denatured in SDS-PAGE sample buffer and resolved on SDS-PAGE. The blot was probed with an anti-mCherry antibody, then stripped and reprobed with an anti-CSP antibody. Detection of the mCherry signal in Triton X-100-treated sporozoites confirmed its presence on the inner membrane. S14 Tr; S14-3XHA-mCherry transgenic parasite. **(C)** The WT GFP and *S14* KO hemolymph sporozoites were allowed to glide for one hour, and the CSP trail left was revealed using a biotin-CSP antibody followed by streptavidin-FITC. All *S14* KO sporozoites were nonmotile. **(D)** Gliding was quantified by counting the CSP trials, and a significant difference was observed (***P<0.0003, one-way ANOVA).

### In silico docking revealed the interaction of S14 with GAP45 and MTIP

The gliding phenotype of the S14 KO and its localization on IMC prompted us to investigate the association of S14 with glideosome-associated proteins. We chose two IMC-localized proteins, MTIP (Bergman *et al*, 2003) and GAP45 (Gaskins *et al*, 2004), for the interaction studies. We started with bioinformatic studies to check the interaction of S14 with MTIP and GAP45. Structural models of MTIP, GAP45 and S14 were obtained and docking was performed (Figure S6). The ClusPro Docking server binding energy of MTIP-GAP45, S14-GAP45, and S14-MTIP were −1216 Kcal/mol, −898.6 Kcal/mol, and −1024.6 Kcal/mol, respectively. Further, we performed the docking using the HDOCK server for the receptor-ligand interface (Figure S7). The set cut-off confidence score was 0.5 and the confidence score for MTIP-GAP45, S14-GAP45 and S14-MTIP interactions were 0.6400, 0.6976 and 0.7235, respectively. These results indicated that MTIP-GAP45 and S14-GAP45 would possibly bind and S14-MTIP are likely to bind. Furthermore, we analyzed protein-protein interaction using HADDOCK server and obtained top Z score clusters were submitted to PDBe_PISA, which revealed interacting residues interface through hydrogen bonding and salt bridge residues. The receptor-ligand interface residues less than 3Å were visualized using Pymol (Figure 5). The respective receptor-ligand interface are given in Tables S3 and S4. The binding energies and residue interface studies indicate that S14 interact with GAP45 and MTIP.

**Figure 5:**
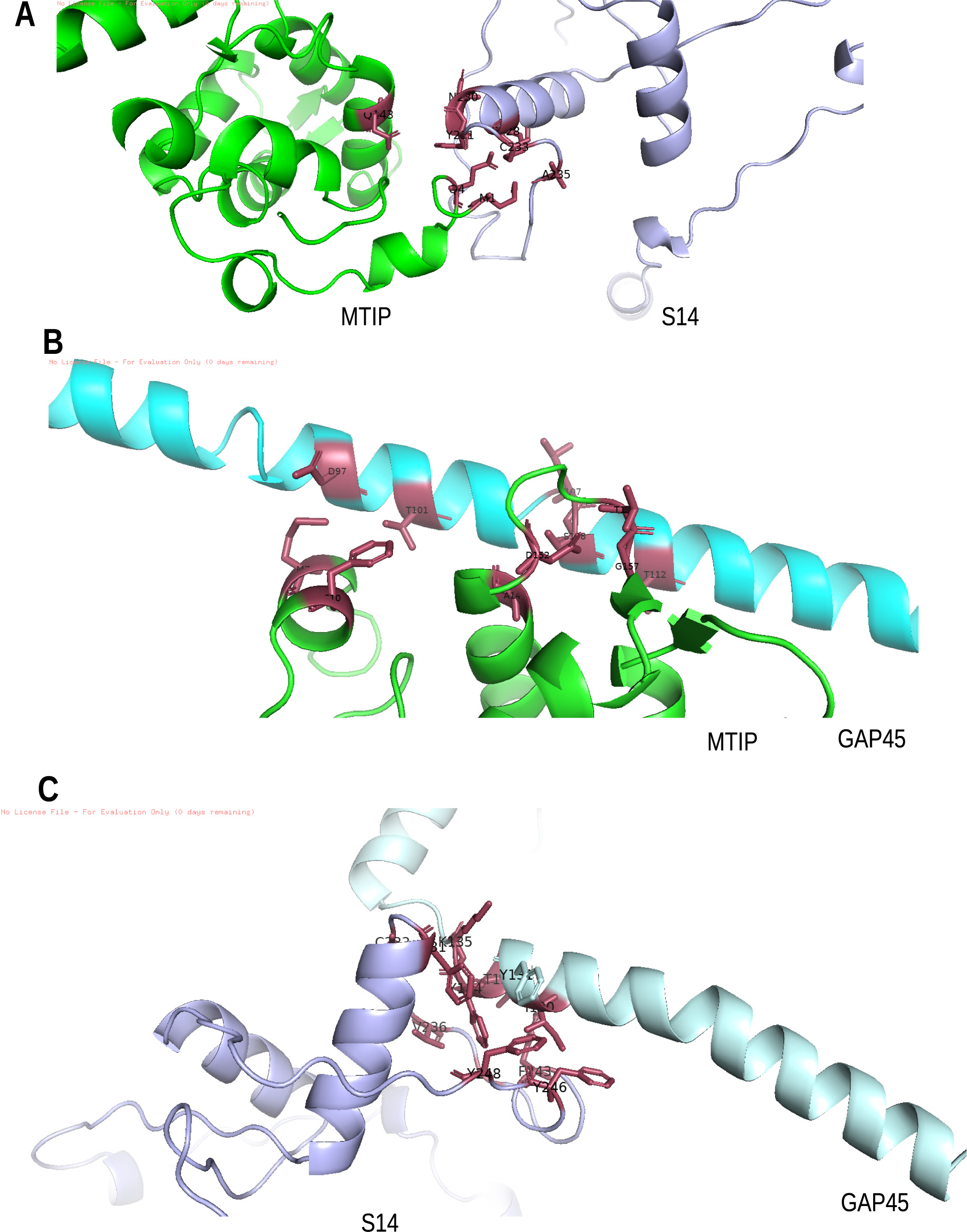
In silico docking studies using HDOCK server showing binding of *P. berghei* S14-MTIP, MTIP-GAP45 and S14-GAP45. A list of receptor-ligand interface are given in Table S4. Images were visualized using Pymol 2.5.4 software. **(A)** S14-MTIP interacting receptor ligand residues (228A CYS-4A GLN, 230A ASN-142A GLU, 231A TYR-4A GLN, 233A CYS-4A GLN, 235A ALA-1A MET). **(B)** MTIP-GAP45 interacting receptor ligand residues (7A MET −97A ASP, 10A PHE-101A THR, 149A ALA-105A LEU, 152A ASP-108A SER, 255A THR-107A LEU, 157A GLY-112A THR). **(C)** S14-GAP45 interacting receptor ligand residues (231A TYR-135A LYS, 233A CYS-134A TYR, 237A VAL-133A THR, 246A TYR-130A ILE, 248A TYR-131A TYR).

### S14 interacts with GAP45 and MTIP in Yeast two-hybrid assay

To further confirm the in-silico interaction results, yeast two-hybrid assay was performed. For this, *P. berghei* S14 was cloned into pAS2, and the MTIP and GAP45 genes were cloned into pGAD-C1 yeast two-hybrid vectors. The plasmid containing the *S14* gene was cotransformed with MTIP or GAP45 genes in the *S. cerevisiae* strain PJ69-4A. Interaction studies were performed using LacZ and HIS3 as reporters and revealed a positive interaction of S14 with the MTIP and GAP45 proteins, as these cells were able to grow on SD-Trp-Leu-His plates containing 10 and 25 mM 3-AT (Figure 6). Furthermore, these cells also gave blue color on plates containing X-gal (Figure 6), indicating a positive interaction of the S14 protein with the MTIP and GAP45 proteins. In contrast, a negative control containing two unrelated proteins could not grow on 3-AT plates and did not give a blue color on the X-gal plate. This data confirmed the interaction of S14 with GAP45 and MTIP.

**Figure 6.**
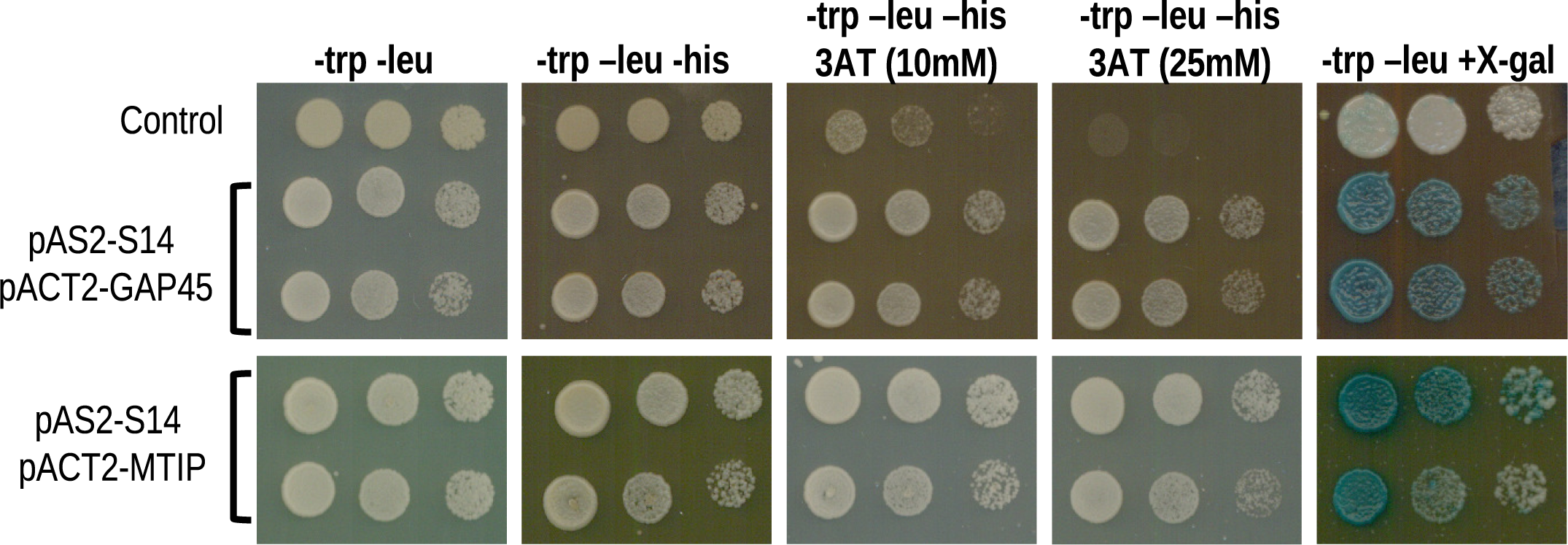
S14 interacts with GAP45 and MTIP in a yeast two-hybrid assay. The S14, MTIP, and GAP45 genes were cloned in a yeast two-hybrid vector. The interaction was analyzed using LacZ as a reporter gene on SD-trp-leu plates containing X-gal and the his3+ marker as a reporter gene on SD-trp-leu plates lacking histidine. 3AT was used to prevent any leaky expression of the his3 marker gene.

### S14 coordinate gliding motility without affecting IMC and surface protein expression and localization

*Plasmodium* sporozoites exhibit a substrate-dependent gliding motility for which surface and IMC proteins are employed. We analyzed the expression and organization of two IMC proteins, MTIP and GAP45, and two surface proteins, CSP and TRAP in *S14* KO sporozoites. Immunostaining revealed an intact IMC and surface organization in *S14* KO sporozoites (Figure 7). This data indicate that S14 performs gliding-specific function and does not affect the organization of IMC and surface proteins.

**Figure 7.**
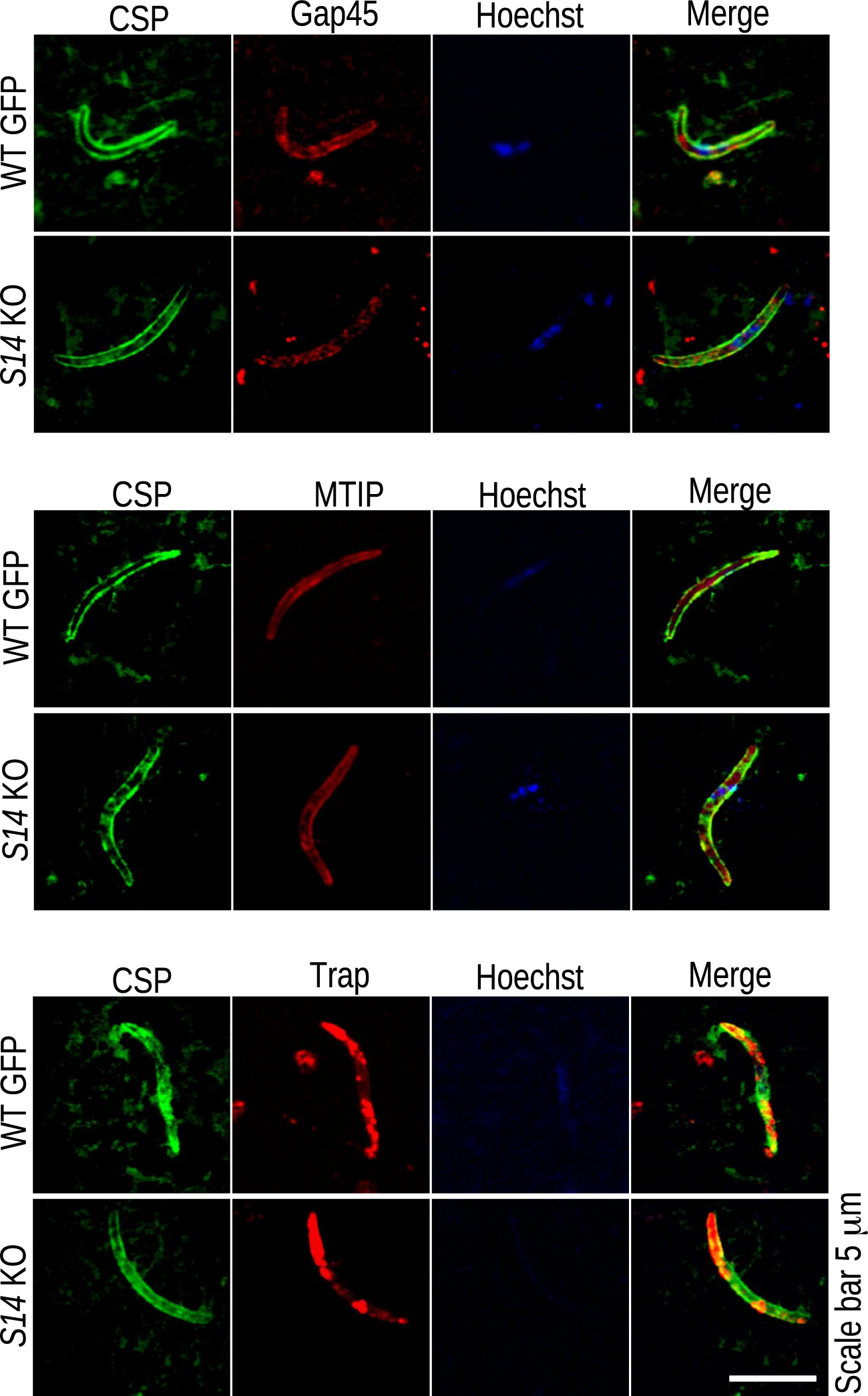
S14 does not affect IMC and surface protein expression and localization. WT GFP and *S14* KO hemolymph sporozoites were spotted on a 12-well slide and air-dried. Immunostaining with anti-GAP45, anti-MTIP, anti-CSP, and anti-TRAP antibodies revealed similar expression and localization in WT GFP and *S14* KO sporozoites.

## Discussion

This study identified a novel *Plasmodium* protein, S14, which lacks signal peptide and transmembrane domains. However, it contains a palmitoylation signal, secreted via the nonclassical pathway and localized to the inner membrane complex. We found that S14 is a gliding-associated protein with a core function in gliding motility and host cell invasion. Deletion of S14 resulted in the accumulation of a higher number of sporozoites in hemolymph, which failed to invade the salivary gland and hepatocytes. The inability of *S14* KO sporozoites to invade the host cell was due to impaired gliding motility. Furthermore, we show that S14 interacts with the glideosome-associated proteins GAP45 and MTIP. Overall, these results indicate a central role for S14 in coordinating gliding motility and invasion of *Plasmodium* sporozoites.

We propose that S14 works with GAP45 and MTIP, facilitating gliding motility. It is possible that S14 connects GAP45 and MTIP or could act entirely independently. Both scenarios are possible because the details of this complex are not known. Similar to S14, several parasite proteins play a role in the gliding motility and invasion of both mosquito and human hosts, such as surface protein TRAP (Sultan et al., 1997) and claudin-like apicomplexan microneme protein (CLAMP) (Loubens *et al*, 2023); however, MAEBL was found to be important for attachment to the salivary gland surface and did not affect sporozoite motility and infectivity to the vertebrate host (Kariu *et al*, 2002). S14 is not a surface protein with an extracellular domain, and its host cell invasion defect was due to impaired gliding. Cellular transmigration and host cell invasion are prerequisites for gliding motility. Several sporozoite proteins, such as SPECT, SPECT2, and PLP1, differ from S14, as they invade the host cell usually with a deficiency in cell traversal (Ishino *et al*, 2004, 2005b; Risco-Castillo *et al*, 2015). Surface proteins such as P36 and P52 show normal gliding motility and cell traversal and play a role in host cell invasion by interacting with host cell receptors (Ishino *et al*, 2005a; Manzoni *et al*, 2017; Arredondo *et al*, 2018).

The parasite gliding machinery consists of the atypical myosin MyoA, MTIP, and ELC1, the glideosome-associated proteins GAP40 and GAP45, and the transmembrane protein GAP50 (Bergman *et al*, 2003; Frénal *et al*, 2010). This MyoA interacts with F-actin, which connects with surface proteins through aldolase (Sultan *et al*, 1997; Jewett & Sibley, 2003; Huynh & Carruthers, 2006). In *T. gondii*, GAP45 plays a vital role in maintaining the close association of the IMC to the plasma membrane (Frénal *et al*, 2010). GAP45 also interacts with GAP50 through its C-terminal region, supporting its function as the anchor of the motor complex in the IMC. We selected GAP45 and MTIP for interaction studies with S14 because MyoA was lost upon the downregulation of MTIP. Furthermore, MTIP was found to be reduced in GAP45 knockdown (Sebastian *et al*, 2012).

Deletion of S14 resulted in the accumulation of a higher number of sporozoites in hemolymph, indicating the dispensable role of S14 during the egress of sporozoites from oocysts. The conditional deletion of GAP45 during *P. falciparum* asexual blood stages revealed its role in the invasion but not in egress. These results indicate that a functional motor complex is not required for egress from RBCs, which plays a critical role in invasion (Perrin *et al*, 2018). GAP40 and GAP50 and members of the GAPM family play critical roles in the biogenesis of IMCs during intracellular replication. Parasites lacking GAP40 or GAP50 start replication but fail to complete it, implicating a structural role in maintaining the stability of the developing IMC during replication (Harding *et al*, 2016). It was shown that IMC is critical for the anchorage and stabilization of the glideosome (Opitz & Soldati, 2002) and is required during the invasion of the host cell (Bargieri *et al*, 2013; Egarter *et al*, 2014; Togbe *et al*, 2008; Meissner *et al*, 2013). We hypothesize that S14 possibly plays a structural role and maintains the stability of IMC required for the activity of motors during gliding and invasion. S14 deletion does not affect GAP45, MTIP, CSP and TRAP expression and localization, suggesting that it performs motor-related functions only. These results indicate that the S14-associated IMC complex possibly exists in sporozoites and coordinates gliding motility (Figure 8); however, the link between S14, GAP45, and MTIP requires further investigation.

**Figure 8.**
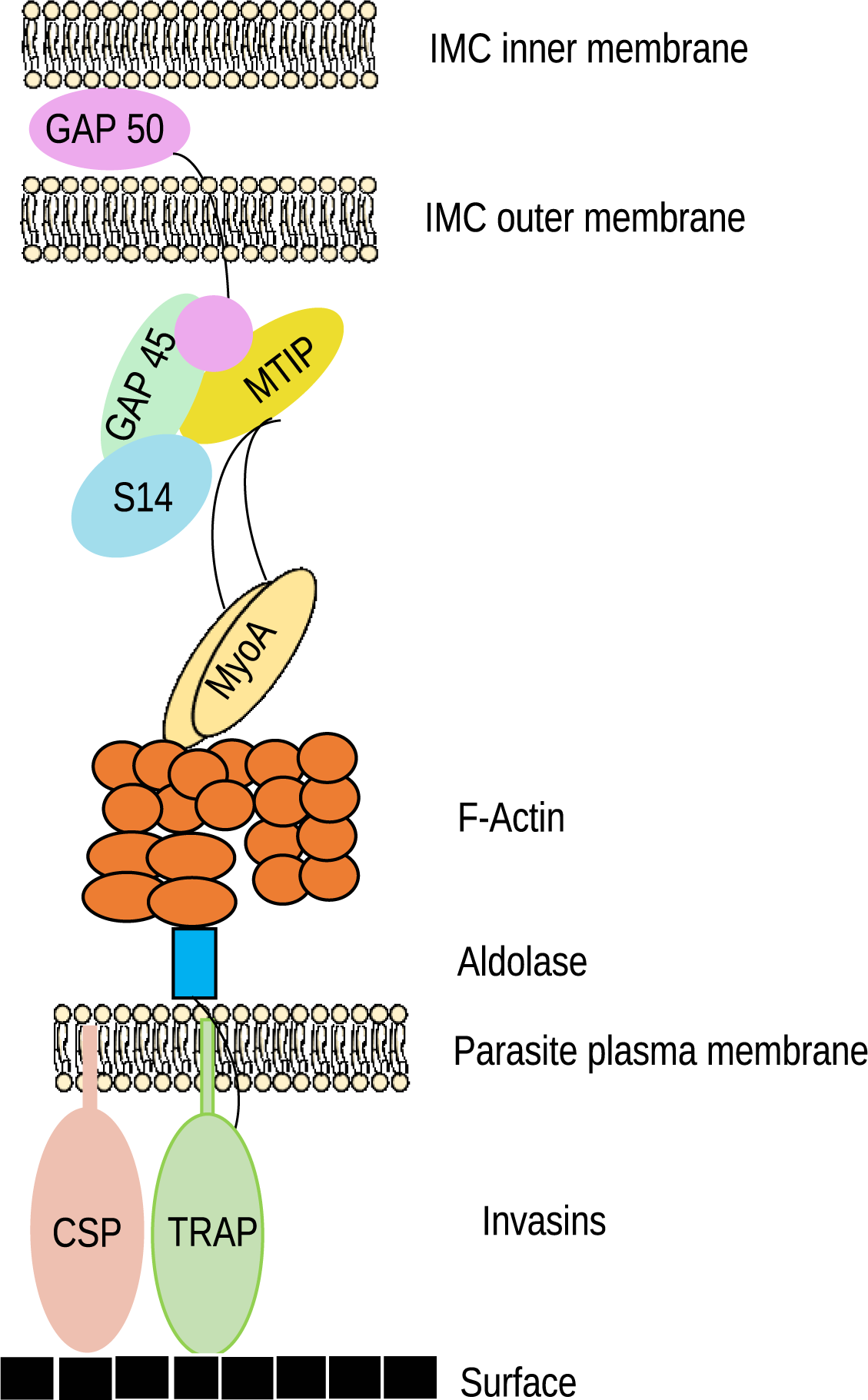
A schematic model of sporozoite membrane with S14. The sporozoite membrane is shown with surface proteins TRAP and CSP. We show that S14 interacts with MTIP and GAP45 and coordinates gliding motility by connecting these two proteins.

## Material and methods

### Parasites, Mosquitos, and Mice

*Plasmodium berghei* ANKA (MRA 311) and *P. berghei* ANKA GFP (MRA 867 507 m6cl1) were obtained from BEI resources, USA. *Anopheles stephensi* mosquitos were reared at 28°C and 80% relative humidity and kept under a 12 h light/dark cycle as previously described (Gupta *et al*, 2020). Swiss albino and C57BL/6 mice were used for parasite infections. All animal procedures were approved by the Institutional Animal Ethics Committee at CSIR-Central Drug Research Institute, India (IAEC/2013/83).

### Protein association prediction for PbS14

Using the “guilt-by-association” principle of prediction, we chose to study the following properties of existing glideosome components along with S14: (1) Classical pathway secretion using the signal peptide (SignalP). (2) Nonclassical pathway secretion (SecretomeP). (3) Presence of transmembrane domains (TMHMM). (4) Presence of a potential palmitoylation site (CSS-Palm). (5) Known associations from the literature. This is a similar association prediction method as employed by the STRING database. These properties were chosen on the rationale that the presence of these signals ascertains protein targeting to different cellular membranes. Associating these characteristics with the known glideosome proteins and their interactions (Boucher & Bosch, 2015), we could select a few glideosome components to check for physical interactions using in-silico studies and yeast two-hybrid assay.

### *S*14 expression analysis by RT‒qPCR

For the absolute quantification of *S14* transcripts, a standard was generated by amplifying a 120 bp fragment within the *S14* ORF (PBANKA_0605900) using primers 1001/1002 (primers are given in Table S5). The amplified product was cloned into the pCR 2.1-TOPO vector. For the normalization of transcripts, Hsp70 was used (Choudhary *et al*, 2019). Total RNA was isolated from blood stage schizonts (Schz), liver stages (LS), midgut (MG), and salivary gland (SG) sporozoites using TRIzol reagent (Takara Bio, Japan) and an RNA isolation kit (Genetix, India) following the manufacturers’ instructions. cDNA was prepared by reverse transcription using a Superscript cDNA synthesis kit. Real-time PCR was carried out using SYBR green reagent (Takara Bio, Japan), and the ratio of transcript numbers of *S14* and *Hsp70* was used to determine the copy number.

### Generation of S14-3XHA-mCherry transgenic parasites

For the endogenous tagging of S14 with 3XHA-mCherry, two fragments, F1 (0.74 kb) and F2 (0.65 kb), were amplified using 1010/1011 and 1005/1006 and cloned into plasmid pBC-3XHA-mCherry-hDHFR at *Xho*I/*Bgl*II and *Not*I/*Asc*I, respectively. The plasmid was linearized using *Xho*I/*Asc*I and transfected into *P. berghei* ANKA schizonts as previously described (Janse *et al*, 2006). Correct 5’ and 3’ site-specific integration was confirmed by diagnostic PCR using primers 1007/1392 and 1215/1008, respectively (primers are given in Table S5). The clonal lines were obtained by limiting dilution of the parasites and analyzed for expression and localization.

### Generation of *S14* KO and complemented parasites

*P. berghei S14* (PBANKA_0605900) was disrupted by double-crossover homologous recombination. For this, two fragments, F3 (0.73 kb) and F4 (0.637 kb), were amplified using *P. berghei* ANKA genomic DNA with primers 1003/1004 and 1005/1006, respectively. The fragments F3 and F4 were cloned sequentially into the pBC-GFP-hDHFR vector at *Xho*I/*Cla*I and *Not*I/*Asc*I, respectively, and finally linearized with *Xho*I/*Asc*I and transfected into *P. berghei* schizonts (Janse *et al*, 2006). The drug-resistant GFP-expressing parasites were confirmed for 5’ and 3’ site-specific integrations using primers 1007/1225 and 1215/1008, respectively (primers are given in Table S5). To generate an *S14* KO complemented parasite line, another *S14* KO parasite line was generated with the hDHFR:yFCU selection cassette. A fragment consisting of the S14 5’UTR, ORF, and 3’UTR was amplified using primers 1003/1006 and transfected into *S14* KO parasites. Parasites containing restored S14 loci were selected by negative selection using a 5-fluorocytosine (5-FC) drug (Sigma, USA) as previously described (Srivastava & Mishra, 2022). The clonal lines were obtained by limiting dilution of the parasites, and the absence of *S14* ORF was confirmed using primers 1010/1011. Furthermore, two clonal lines were also confirmed by Southern blotting as described previously (Narwal *et al*, 2022). Fragment F3 was used as a probe to detect the band in Southern blot.

### Phenotypic characterization of *S14* KO parasites

The *S14* KO clonal lines were first analyzed for asexual blood-stage propagation, and for this, 200 µl of iRBCs with 0.2% parasitemia was intravenously injected into a group of mice. Parasitemia was monitored daily by Giemsa staining of blood smears. Next, we initiated infection with KO parasites in *Anopheles stephensi* mosquitos as previously described (Narwal *et al*, 2022). On days 14 and 19, midgut and salivary glands were observed for infection, and sporozoite numbers were counted. The hemolymph sporozoites were collected and counted as previously described (Mastan *et al*, 2017).

### In vivo infectivity

To determine the in vivo infectivity of KO sporozoites, C57BL/6 mice were either infected by mosquito bite or by intravenously injecting hemolymph sporozoites. For the bite experiment, ten mosquitos per cage were used. The appearance of parasites in the blood was observed by making Giemsa-stained blood smears.

### In vitro infectivity of sporozoites

The in vitro infectivity of sporozoites was tested by infecting HepG2 cells as previously described (Narwal *et al*, 2022). Fifty thousand cells/well were seeded in 48-well plates containing sterilized coverslips pretreated with collagen. Hemolymph sporozoites (10,000 sporozoites/well for the invasion assay or 5,000 sporozoites/well for EEF development) were added to the HepG2 monolayers, and the plate was centrifuged at 310 g for 4 min and incubated at 37°C in a CO2 incubator. The culture was fixed using 4% PFA at 1 hpi for the invasion assay and 40 hpi for the EEF development assay.

### Transformation of sporozoites into early EEFs without host cells

WT GFP and *S14* KO hemolymph sporozoites were incubated in a medium containing DMEM with 2 mM L-glutamine, 4.5 g/liter glucose, and supplemented with 10% FBS (Sigma, USA), 500 U/ml penicillin‒streptomycin, (Thermo Fisher Scientific, USA) and 1.25 μl/ml fungizone as previously described (Kaiser *et al*, 2003). The sporozoites were incubated at 37°C in a CO2 incubator for 4 hrs and then fixed using 4% PFA.

### Generation of anti-MTIP and anti-GAP45 antibodies

Affinity-purified polyclonal rabbit antibodies against *P. berghei* MTIP and GAP45 were developed by GenScript Inc., Piscataway, NJ, against the peptide sequences CVNKDDRKIYFDEKS and CHKYENDSDKLETGS, respectively.

### Triton X-100 membrane extraction

Sporozoites (3 ×10^4^) were collected and treated with 1.0% Triton X-100 diluted in PBS and incubated on ice for 30 min as previously described (Bergman *et al*, 2003). After incubation, sporozoites were spun at 13,800 × g for 20 min at 4°C. Both treated and untreated sporozoites were washed three times with PBS and fixed with 2% paraformaldehyde diluted in PBS or resuspended.

### Sporozoite gliding motility assay

To quantify sporozoite gliding motility, a glass eight-well chamber slide was coated with 10 μg/ml anti-CSP antibody in PBS overnight, and the assay was performed as described previously (Stewart & Vanderberg, 1988). Hemolymph sporozoites collected in 3% BSA/DMEM were added at 5,000/well and incubated for 1 h at 37°C in a CO2 incubator. After incubation, sporozoites were fixed with 4% PFA, blocked using 3% BSA/PBS, and incubated with biotin-labeled anti-CSP antibody, and signals were revealed using streptavidin-FITC (Invitrogen, USA). Trails associated with sporozoites were counted using a Nikon 80i fluorescence microscope.

### Immunofluorescence assay

Fixed sporozoites were washed with PBS, permeabilized using 0.1% Triton X-100 for 15 min at room temperature, and then blocked with 1% BSA/PBS for 1 hr at room temperature. Further sporozoites were incubated with anti-mCherry developed in rabbit (Novus Biologicals, USA), anti-CSP mouse monoclonal (Yoshida *et al*, 1980), anti-MTIP, anti-GAP45 or anti-TRAP (Mishra *et al*, 2023) antibodies. The signals were revealed using Alexa Fluor 594-conjugated or Alexa Fluor 488-conjugated antibodies (diluted 1:1,000; Invitrogen, USA). For the staining of EEFs, fixed cultures were washed with PBS, and staining with anti-UIS4 antibody (Mueller *et al*, 2005) was performed as previously described (Narwal *et al*, 2022). Nuclei were stained with Hoechst 33342 (Sigma‒Aldrich, USA) and mounted using Prolong diamond anti-fade reagent (Invitrogen, USA). The images were acquired using a confocal laser scanning microscope with a UPlanSAPO 100x/1.4 oil immersion objective (Olympus BX61WI) at 4X magnification.

### Western blot analysis

Sporozoites were pelleted by centrifuging at 14,000 rpm for 4 min and resuspended in Laemmli buffer. Immunoblotting was performed as previously described (Narwal *et al*, 2022). Briefly, samples were resolved by SDS-PAGE, transferred to a nitrocellulose membrane (Bio-Rad, USA), and blocked with 1% BSA. The blot was incubated with an anti-mCherry (diluted 1:1,000, Novus Biologicals, USA) or anti-CSP (Yoshida *et al*, 1980) antibodies. The membrane was washed and incubated with HRP-conjugated anti-rabbit or anti-mouse IgG (diluted 1:5,000, Amersham Biosciences, United Kingdom). The signals were detected using ECL chemiluminescent substrate (Thermo Scientific, USA) in a ChemiDoc XRS+ System (Bio-Rad, USA).

### Bioinformatic approaches for interrogating protein-protein interactions

To check the interaction of S14 (PBANKA_0605900) with MTIP (PBANKA_1459500) and GAP45 (PBANKA_1437600), in-silico docking was performed. The PDB structures of S14, MTIP and GAP45 are not available, therefore, to obtain their model structure, amino acids sequences were submitted to trRosetta server (https://yanglab.nankai.edu.cn/trRosetta). Obtained structural models were validated using SAVE webserver (https://saves.mbi.ucla.edu/) for Quality factors and Ramachandran plots acceptibility. Models with quality factor above 95% were further processed for the protein-protein interaction studies. ClusPro server (https://cluspro.bu.edu/home.php) was used for initial protein-protein docking studies to predict MTIP-GAP45, MTIP-S14 and S14-GAP45 interaction according to their binding energy. Further, PDB files of the models were submitted to the HDOCK server (http://hdock.phys.hust.edu.cn) for their interaction studies using default parameters. The model with the highest docking score was further used for visualization by Pymol 2.5.4 software (http://www.pymol.org/pymol) (Schrödinger & DeLano).

### Yeast two-hybrid interaction studies

The *P. berghei* S14 gene was amplified using primers 2058/2059 and cloned into the Gal4 binding domain-containing vector pAS2 for yeast two-hybrid interaction studies. The MTIP and GAP45 genes were amplified using primer pairs 2060/2061 and 2062/2063, respectively, and cloned into the pGAD-C1 vector in frame with the Gal4 activation domain. These plasmids were cotransformed into *S. cerevisiae* strain PJ69-4A (Clontech, USA), which contains the lacZ gene from *E. coli* encoding β-galactosidase and the HIS3 selectable marker as reporter genes. Interaction studies were performed on SD-leu-trp plates containing X-gal or SD-leu-trp-his plates containing 3-aminotriazole (3-AT). The appearance of blue color on X-gal plates and growth on plates containing 10 and 25 mM 3-AT in the absence of histidine indicates a positive interaction.

### Statistical analysis

Statistical analysis was done using two-tailed, unpaired Student’s t test or one-way ANOVA in GraphPad Prism software.

## Data and Materials Availability

All data are available within this manuscript, and raw data are available from the corresponding author upon reasonable request. Materials generated in this study are available from the corresponding author on request.

## Acknowledgments

We thank Dr. Pratik Narain Srivastava for performing initial bioinformatic studies. We thank BEI Resources, USA, for the parasite strains and plasmids and Dr. Kota Arun Kumar (University of Hyderabad, India) for the pBC-3XHA-mCherry-hDHFR vector. We thank Dr. Photini Sinnis (Johns Hopkins University, USA) for the anti-UIS4 antibody. We acknowledge the THUNDER (BSC0102) and MOES (GAP0118) Intravital and Confocal microscopy facility of CSIR-CDRI. The Council of Scientific and Industrial Research, University Grants Commission, and Indian Council of Medical Research, Government of India research fellowships supported AG, AV, N, RG, SG and SKN. The Ramalingaswami Fellowship grant supported the work (BT/RLF/Re-entry/20/2012). This manuscript is CDRI communication no. 144/2023/SM.

## Author contributions

AG, AV and SM conceived the idea, designed and performed the experiments, and analyzed the data. SKN and RG performed the experiments. N completed the in-silico protein-protein interaction studies. SG and SA performed the yeast two-hybrid interaction studies. SM wrote the manuscript, and all the authors have read and approved the manuscript.

## References

Arredondo SA, Swearingen KE, Martinson T & Steel R (2018) The Micronemal Plasmodium Proteins P36 and P52 Act in Concert to Establish the Compartment Within Infected Hepatocytes. 8: 1–17

Bargieri DY, Andenmatten N, Lagal V, Thiberge S, Whitelaw JA, Tardieux I, Meissner M & Ménard R (2013) Apical membrane antigen 1 mediates apicomplexan parasite attachment but is dispensable for host cell invasion. Nat Commun 4

Baum J, Gilberger TW, Frischknecht F & Meissner M (2008) Host-cell invasion by malaria parasites: insights from Plasmodium and Toxoplasma. Trends Parasitol 24: 557–563

Baum J, Richard D, Healer J, Rug M, Krnajski Z, Gilberger TW, Green JL, Holder AA & Cowman AF (2006) A conserved molecular motor drives cell invasion and gliding motility across malaria life cycle stages and other apicomplexan parasites. J Biol Chem 281: 5197–5208

Bergman LW, Kaiser K, Fujioka H, Coppens I, Daly TM, Fox S, Matuschewski K, Nussenweig V & Kappe SHI (2003) Myosin A tail domain interacting protein (MTIP) localizes to the inner membrane complex of Plasmodium sporozoites. J Cell Sci 116: 39–49

Boucher LE & Bosch J (2015) The apicomplexan glideosome and adhesins - Structures and function. J Struct Biol 190: 93–114

Buscaglia CA, Coppens I, Hol WGJ & Nussenzweig V (2003) Sites of interaction between aldolase and thrombospondin-related anonymous protein in plasmodium. Mol Biol Cell 14: 4947–4957

Choudhary HH, Gupta R & Mishra S (2019) PKAc is not required for the preerythrocytic stages of Plasmodium berghei. 2: 1–11

Combe A, Moreira C, Ackerman S, Thiberge S, Templeton TJ & Ménard R (2009) TREP, a novel protein necessary for gliding motility of the malaria sporozoite. Int J Parasitol 39: 489–496

Daher W & Soldati-Favre D (2009) Mechanisms controlling glideosome function in apicomplexans. Curr Opin Microbiol 12: 408–414

Douglas RG, Amino R, Sinnis P & Frischknecht F (2015) Active migration and passive transport of malaria parasites. Trends Parasitol 31: 357–362

Egarter S, Andenmatten N, Jackson AJ, Whitelaw JA, Pall G, Black JA, Ferguson DJP, Tardieux I, Mogilner A & Meissner M (2014) The toxoplasma Acto-MyoA motor complex is important but not essential for gliding motility and host cell invasion. PLoS One 9: e91819

Frénal K, Dubremetz JF, Lebrun M & Soldati-Favre D (2017) Gliding motility powers invasion and egress in Apicomplexa. Nat Rev Microbiol 15: 645–660

Frénal K, Polonais V, Marq JB, Stratmann R, Limenitakis J & Soldati-Favre D (2010) Functional dissection of the apicomplexan glideosome molecular architecture. Cell Host Microbe 8: 343–357

Gaskins E, Gilk S, DeVore N, Mann T, Ward G & Beckers C (2004) Identification of the membrane receptor of a class XIV myosin in Toxoplasma gondii. J Cell Biol 165: 383– 393

Giovannini D, Späth S, Lacroix C, Perazzi A, Bargieri D, Lagal V, Lebugle C, Combe A, Thiberge S, Baldacci P, et al (2011) Independent roles of apical membrane antigen 1 and rhoptry neck proteins during host cell invasion by apicomplexa. Cell Host Microbe 10: 591–602

Gould SB, Kraft LGK, Van Dooren GG, Goodman CD, Ford KL, Cassin AM, Bacic A, McFadden GI & Waller RF (2011) Ciliate pellicular proteome identifies novel protein families with characteristic repeat motifs that are common to alveolates. Mol Biol Evol 28: 1319–1331

Gupta R, Mishra A, Choudhary HH, Narwal SK, Nayak B, Srivastava PN & Mishra S (2020) Secreted protein with altered thrombospondin repeat (SPATR) is essential for asexual blood stages but not required for hepatocyte invasion by the malaria parasite Plasmodium berghei. Mol Microbiol 113: 478–491

Harding CR, Egarter S, Gow M, Jiménez-Ruiz E, Ferguson DJP & Meissner M (2016) Gliding Associated Proteins Play Essential Roles during the Formation of the Inner Membrane Complex of Toxoplasma gondii. PLoS Pathog 12: e1005403

Heintzelman MB (2015) Gliding motility in apicomplexan parasites. Semin Cell Dev Biol 46: 135–142

Heiss K, Nie H, Kumar S, Daly TM, Bergman LW & Matuschewski K (2008) Functional characterization of a redundant Plasmodium TRAP family invasin, TRAP-like protein, by aldolase binding and a genetic complementation test. Eukaryot Cell 7: 1062–1070

Huynh M & Carruthers VB (2006) Toxoplasma MIC2 Is a Major Determinant of Invasion and Virulence. 2

Ishino T, Chinzei Y & Yuda M (2005a) Two proteins with 6-cys motifs are required for malarial parasites to commit to infection of the hepatocyte. Mol Microbiol 58: 1264– 1275

Ishino T, Chinzei Y & Yuda M (2005b) A Plasmodium sporozoite protein with a membrane attack complex domain is required for breaching the liver sinusoidal cell layer prior to hepatocyte infection. Cell Microbiol 7: 199–208

Ishino T, Yano K, Chinzei Y & Yuda M (2004) Cell-passage activity is required for the malarial parasite to cross the liver sinusoidal cell layer. PLoS Biol 2: E4

Janse CJ, Ramesar J & Waters AP (2006) High-efficiency transfection and drug selection of genetically transformed blood stages of the rodent malaria parasite Plasmodium berghei. Nat Protoc 1: 346–356

Jewett TJ & Sibley LD (2003) Aldolase forms a bridge between cell surface adhesins and the actin cytoskeleton in apicomplexan parasites. Mol Cell 11: 885–894

Kaiser K, Camargo N & Kappe SHI (2003) Transformation of sporozoites into early exoerythrocytic malaria parasites does not require host cells. J Exp Med 197: 1045–1050

Kaiser K, Matuschewski K, Camargo N, Ross J & Kappe SHI (2004) Differential transcriptome profiling identifies Plasmodium genes encoding pre-erythrocytic stage-specific proteins. Mol Microbiol 51: 1221–1232

Kariu T, Yuda M, Yano K & Chinzei Y (2002) MAEBL is essential for malarial sporozoite infection of the mosquito salivary gland. J Exp Med 195: 1317–1323

Kehrer J, Singer M, Lemgruber L, Silva PAGC, Frischknecht F & Mair GR (2016) A Putative Small Solute Transporter Is Responsible for the Secretion of G377 and TRAP-Containing Secretory Vesicles during Plasmodium Gamete Egress and Sporozoite Motility. 1–25

King CA (1988) Cell motility of sporozoan protozoa. Parasitol Today 4: 315–319

Klug D & Frischknecht F (2017) Motility precedes egress of malaria parasites from oocysts. Elife 6: 1–32

Kono M, Prusty D, Parkinson J & Gilberger TW (2013) The apicomplexan inner membrane complex. Front Biosci (Landmark Ed 18: 982–992

Labaied M, Camargo N & Kappe SHII (2007) Depletion of the Plasmodium berghei thrombospondin-related sporozoite protein reveals a role in host cell entry by sporozoites. Mol Biochem Parasitol 153: 158–166

Loubens M, Marinach C, Paquereau C & Hamada S (2023) The claudin-like apicomplexan microneme protein is required for gliding motility and infectivity of Plasmodium sporozoites. 1–24

Manzoni G, Marinach C, Topçu S, Briquet S, Grand M, Tolle M, Gransagne M, Lescar J, Andolina C, Franetich JF, et al (2017) Plasmodium P36 determines host cell receptor usage during sporozoite invasion. Elife 6: 1–24

Mastan BS, Narwal SK, Dey S, Kumar KA & Mishra S (2017) Plasmodium berghei plasmepsin VIII is essential for sporozoite gliding motility. Int J Parasitol 47: 239–245

Meissner M, Ferguson DJP & Frischknecht F (2013) Invasion factors of apicomplexan parasites: essential or redundant? Curr Opin Microbiol 16: 438–444

Mikolajczak SA, Silva-Rivera H, Peng X, Tarun AS, Camargo N, Jacobs-Lorena V, Daly TM, Bergman LW, de la Vega P, Williams J, et al (2008) Distinct Malaria Parasite Sporozoites Reveal Transcriptional Changes That Cause Differential Tissue Infection Competence in the Mosquito Vector and Mammalian Host. Mol Cell Biol 28: 6196– 6207

Mishra A, Srivastava PN, H SA & Mishra S (2023) Autophagy protein Atg7 is essential and druggable for maintaining malaria parasite cellular homeostasis and organelle biogenesis. bioRxiv: 2023.08.16.553492

Mueller AK, Camargo N, Kaiser K, Andorfer C, Frevert U, Matuschewski K & Kappe SHI (2005) Plasmodium liver stage developmental arrest by depletion of a protein at the parasite-host interface. Proc Natl Acad Sci U S A 102: 3022–3027

Narwal SK, Nayak B, Mehra P & Mishra S (2022) Protein kinase 9 is not required for completion of the Plasmodium berghei life cycle. Microbiol Res 260: 127051

Opitz C & Soldati D (2002) ‘The glideosome’: a dynamic complex powering gliding motion and host cell invasion by Toxoplasma gondii. Mol Microbiol 45: 597–604

Perrin AJ, Collins CR, Russell MRG, Collinson LM, Baker DA & Blackman MJ (2018) The actinomyosin motor drives malaria parasite red blood cell invasion but not egress. MBio 9: 1–13

Poulin B, Patzewitz E-M, Brady D, Silvie O, Wright MH, Ferguson DJP, Wall RJ, Whipple S, Guttery DS, Tate EW, et al (2013) Unique apicomplexan IMC sub-compartment proteins are early markers for apical polarity in the malaria parasite. Biol Open 2: 1160– 1170

Prudêncio M, Rodriguez A & Mota MM (2006) The silent path to thousands of merozoites: The Plasmodium liver stage. Nat Rev Microbiol 4: 849–856

Rees-Channer RR, Martin SR, Green JL, Bowyer PW, Grainger M, Molloy JE & Holder AA (2006) Dual acylation of the 45 kDa gliding-associated protein (GAP45) in Plasmodium falciparum merozoites. Mol Biochem Parasitol 149: 113–116

Rénia L, Miltgen F, Charoenvit Y, Ponnudurai T, Verhave JP, Collins WE & Mazier D (1988) Malaria sporozoite penetration A new approach by double staining. J Immunol Methods 112: 201–205

Ripp J, Kehrer J, Smyrnakou X, Tisch N, Tavares J, Amino R, Ruiz de Almodovar C & Frischknecht F (2021) Malaria parasites differentially sense environmental elasticity during transmission. EMBO Mol Med 13: 1–10

Risco-Castillo V, Topçu S, Marinach C, Manzoni G, Bigorgne AE, Briquet S, Baudin X, Lebrun M, Dubremetz J-F & Silvie O (2015) Malaria Sporozoites Traverse Host Cells within Transient Vacuoles. Cell Host Microbe 18: 593–603

Saenz FE, Balu B, Smith J, Mendonca SR & Adams JH (2008) The transmembrane isoform of Plasmodium falciparum MAEBL is essential for the invasion of Anopheles salivary glands. PLoS One 3

Schrödinger LLC & DeLano W PyMOL. [PREPRINT]

Sebastian S, Brochet M, Collins MO, Schwach F, Jones ML, Goulding D, Rayner JC, Choudhary JS & Billker O (2012) A Plasmodium calcium-dependent protein kinase controls zygote development and transmission by translationally activating repressed mRNAs. Cell Host Microbe 12: 9–19

Srivastava PN & Mishra S (2022) Disrupting a Plasmodium berghei putative phospholipase impairs efficient egress of merosomes. Int J Parasitol 52: 547–558

Steinbuechel M & Matuschewski K (2009) Role for the Plasmodium sporozoite-specific transmembrane protein S6 in parasite motility and efficient malaria transmission. Cell Microbiol 11: 279–288

Stewart MJ & Vanderberg JP (1988) Malaria sporozoites leave behind trails of circumsporozoite protein during gliding motility. J Protozool 35: 389–393

Sultan AA, Thathy V, Frevert U, Robson KJ, Crisanti A, Nussenzweig V, Nussenzweig RS & Ménard R (1997) TRAP is necessary for gliding motility and infectivity of plasmodium sporozoites. Cell 90: 511–522

Talman AM, Lacroix C, Marques SR, Blagborough AM, Carzaniga R, Ménard R & Sinden RE (2011) PbGEST mediates malaria transmission to both mosquito and vertebrate host. 82: 462–474

Togbe D, de Sousa PL, Fauconnier M, Boissay V, Fick L, Scheu S, Pfeffer K, Menard R, Grau GE, Doan BT, et al (2008) Both functional LTβ receptor and TNF receptor 2 are required for the development of experimental cerebral malaria. PLoS One 3

Yeoman JA, Hanssen E, Maier AG, Klonis N, Maco B, Baum J, Turnbull L, Whitchurch CB, Dixon MWA & Tilley L (2011) Tracking Glideosome-Associated Protein 50 Reveals the Development and Organization of the Inner Membrane Complex of. 10: 556–564

Yoshida N, Nussenzweig RS, Potocnjak P, Nussenzweig V & Aikawa M (1980) Hybridoma Produces Protective Antibodies Directed against the Sporozoite Stage of Malaria Parasite. Science (80-) 207: 71–73

